# Speciation without gene-flow in hybridising deer

**DOI:** 10.1101/2022.04.20.488928

**Authors:** Camille Kessler, Eric Wootton, Aaron B.A. Shafer

## Abstract

Under the ecological speciation model, divergent selection acts on ecological differences between populations, gradually creating barriers to gene flow and ultimately leading to reproductive isolation. Hybridisation is part of this continuum and can both promote and inhibit the speciation process. Here, we used white-tailed (*Odocoileus virginianus*) and mule deer (*O. hemionus*) to investigate patterns of speciation in hybridising sister species. We quantified genome-wide historical introgression and performed genome scans to look for signatures of four different selection scenarios. Despite ample modern evidence of hybridisation, we found negligible patterns of ancestral introgression and no signatures of divergence with gene flow, rather localised patterns of allopatric and balancing selection were detected across the genome. Genes under balancing selection were related to immunity, MHC and sensory perception of smell, the latter of which is consistent with deer biology. The deficiency of historical gene-flow suggests that white-tailed and mule deer were spatially separated during the glaciation cycles of the Pleistocene and genome wide differentiation accrued via genetic drift. Dobzhansky-Muller incompatibilities and selection against hybrids are hypothesised to be acting, and diversity correlations to recombination rates suggests these sister species are far along the speciation continuum.

## Introduction

Hybridisation is a widespread phenomenon that occurs at variable rates (Adavoudi & Pilot, 2021; Iacolina et al., 2019; Mallet, 2005; Ragavan et al., 2017; Taylor & Larson, 2019). The prevalence of hybridisation suggests speciation follows a continuum as opposed to the more discrete and allopatric view introduced by Mayr (1942) (Mallet, 2005; Stankowski & Ravinet, 2021). However, both ideas are not incompatible as the speciation continuum ultimately leads to reproductive isolation with hybridisation being a natural outcome (Stankowski & Ravinet, 2021). Under the ecological speciation model (sensu Darwin, 1859), hybrids are common and can even facilitate speciation (Nosil, 2012). Ecological speciation works when ecological differences lead to divergent selection between populations; the underlying genetic mechanisms (i.e. causative loci) gradually create barriers to gene flow that accumulate which leads to reproductive isolation (Nosil, 2012; Rundle & Nosil, 2005; Shafer & Wolf, 2013).

The genetic mechanisms involved in ecological speciation can take the form of barrier loci which act as a restraint to gene flow in different ways: for example, such loci can be under divergent selection, involved in mate choice, contribute to assortative mating, or reduce hybrid fitness (Abbott et al., 2013; Ravinet et al., 2017). Divergent selection should produce genomic islands in this context, also referred to as genomic islands of speciation, that are regions where the differentiation (e.g. F_ST_) is higher than the neutral genomic background (Campbell et al., 2018; Ravinet et al., 2017). Such loci and islands have been observed in several species underpinning a wide array of speciation and divergence processes (Lavretsky et al., 2019; Marques et al., 2019; Momigliano et al., 2017; Poelstra et al., 2014). Scanning the genome for just F_ST_ peaks when speciation with gene flow is ongoing, however, is problematic as F_ST_ is dependent on genetic diversity (*π*), and other mechanisms can create similar F_ST_ profiles such as genetic drift, global adaptation, or simply reduced genetic diversity due to background selection (Booker et al., 2021; Burri, 2017; Cruickshank & Hahn, 2014; Ravinet et al., 2017). Joint genome scans including F_ST_, d_xy_ and *π* facilitate a better depiction of the processes of selection and ecological speciation, particularly the use of d_xy_ which is not influenced by current levels of diversity (Campbell et al., 2018; Cruickshank & Hahn, 2014; Ravinet et al., 2017; Shang et al., 2021). Shang et al. (2021) recently conceptualised the expected patterns F_ST_, d_xy_ and *π* under four main modes of selection: divergence with gene flow, allopatric selection (i.e., positive selection post species split), balancing selection and background selection. Using pairs of *Populus* species along the speciation continuum, they investigated the genomic landscape to test clear predictions on the behaviour of F_ST_, d_xy_ and *π*. Including correlations of *π* and F_ST_ to recombination rate (ρ) can help place a species on the continuum, with for example the correlation between F_ST_ and ρ is expected to become stronger with increasing divergence (Burri, 2017; Shang et al., 2021). This approach ultimately allows for disentangling selection patterns from a neutral background based on those statistics, and helps decipher the evolutionary history of species.

White-tailed (*Odocoileus virginianus*; WTD) and mule deer (*O. hemionus*; MD) are abundant in North America with similar morphology, activity patterns and life-history traits, but they differ in several ecological aspects (Douzery & Randi, 1997; Gilbert et al., 2006; Pitra et al., 2004; Brunjes et al., 2006; Berry et al., 2019). Mule deer favour food availability whereas WTD prioritise security and thermoregulation (Whittaker & Lindzey, 2004); consequently, WTD prefer habitats at lower altitude and with denser visual cover than MD that favour open areas at higher elevation (Anthony & Smith, 1977; Brunjes et al., 2006). White-tailed deer and MD also differ in their sociality and associated predator response. Mule deer live in large cohesive groups including both sexes, and, as a group, adopt an aggressive behaviour in presence of predators, whereas WTD live in smaller female-biased groups and flee in response to predators (Lingle, 2001, 2003). Both species represent a high economic value in North America as hunting-related activities generate billions of dollars annually (Cambronne, 2013), and both species are an important cultural component of Indigenous communities (Adams & Hamilton, 2011; Peres & Altman, 2018).

Despite a species divergence date estimated at ~3.13 mya (Wright et al., 2022), WTD and MD hybridise in areas of sympatry with estimated hybridisation rates ranging from 1 to 19% depending on the region (Carr & Hughes, 1993; Combe et al., 2021; Hornbeck & Mahoney, 2000; Russell et al., 2021). Based on divergent species distributions (Fig. 1A) and likely separate refugia during glacial events (Greenslade, 1998), both species likely spent considerable time in allopatry. Recent hybrid zone analyses mainly found backcrossed individuals rather than F1 hybrids (Combe et al., 2021; Russell et al., 2021), and an interspecies F_ST_ up to 0.4 (Combe et al., 2021). It has also been shown that gene flow appears restricted but bidirectional, and there are signs of introgression at both mitochondrial and nuclear levels (Bradley et al., 2003; Carr et al., 1986; Carr & Hughes, 1993; Cathey et al., 1998; Cronin, 1991; Derr, 1991; Russell et al., 2019), such that some MD acquired WTD mitochondrial DNA around ~1.32 Mya (Wright et al., 2022). This pattern would suggest ancestral hybridisation and gene flow has taken place. Given their clear behavioural differences, and their documented hybridisation, white-tailed and mule deer do not present the reproductive isolation required by Mayr’s biological species concept (Mayr, 1942). Here, we hypothesised that WTD and MD have evolved via ecological speciation, and more specifically a speciation with gene flow scenario. To test this hypothesis, we quantified genome-wide introgression and past admixture events and performed genome scans to look for signatures of four different divergence scenarios. We expected to find genetic signs of divergent selection, such as speciation islands, consistent with patterns of divergence with gene flow (DwGF), and a stronger F_ST_ - ρ correlation between species compared to within. We also predicted higher rates of admixture and historical introgression in areas of sympatry.

**Figure 1:**
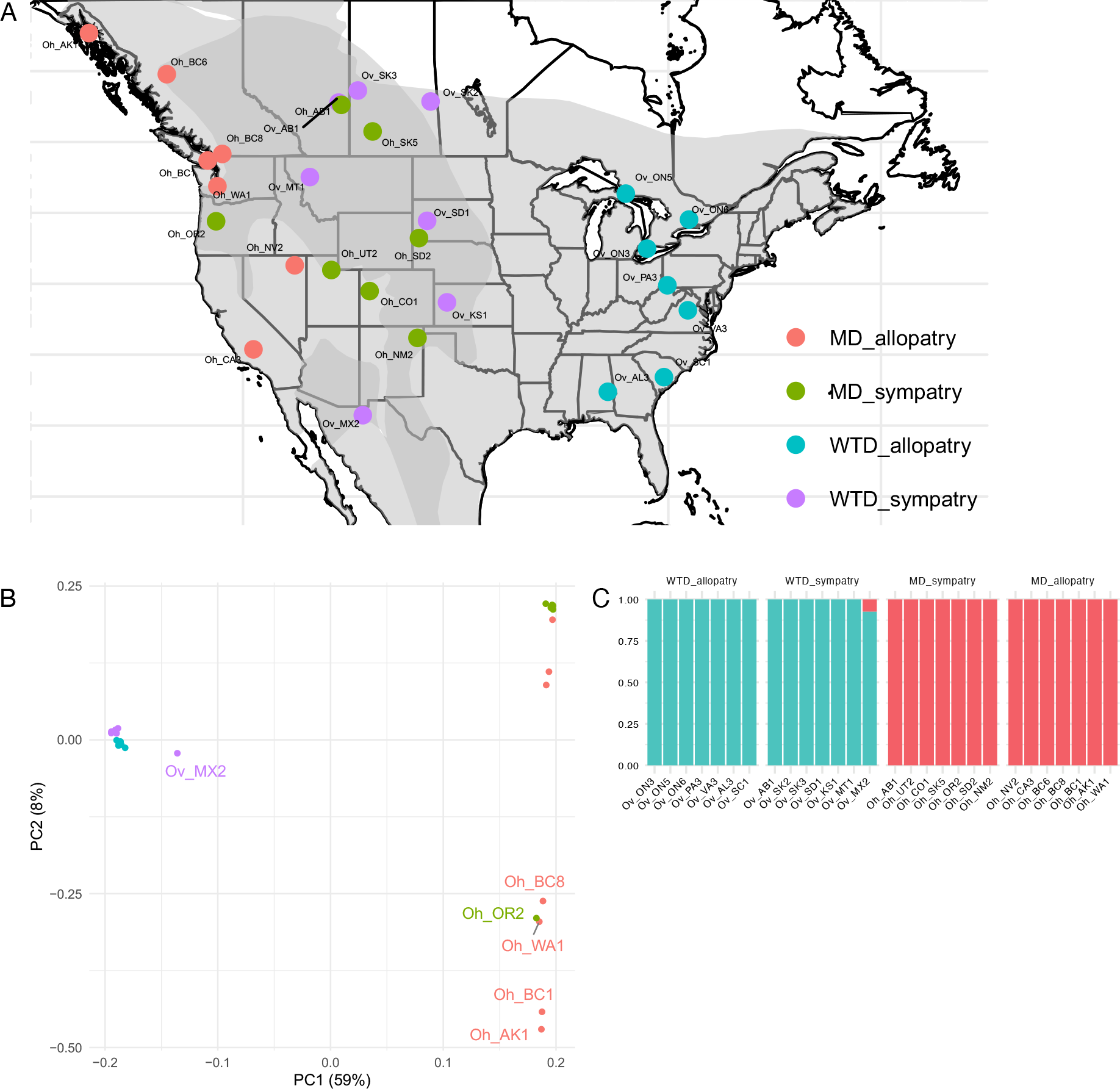
Deer individuals in the study. (A) Sampling locations, colours represent the different groups: WTD allopatry (WTDa, blue), WTD sympatry (WTDs, violet), MD sympatry (MDs, green), MD allopatry (MDa, orange). Areas of sympatry are coloured in dark grey and areas of allopatry in light grey, from the IUCN range data for both species (IUCN, 2015a, 2015b). (B) Principal component analysis performed in PCAngsd. (C) Admixture proportions for K = 2 calculated in NGSAdmix.

## Materials and Methods

### Sampling & Sequencing

We obtained tissue samples from harvested deer collected across the range of WTD & MD including areas of sympatry and allopatry (Fig. 1A, Table 1). These areas were determined from NatureServe records and adjusted with IUCN range data (IUCN, 2015a, 2015b; NatureServe, 2021a, 2021b). Specifically, the state of Washington and the province of British Columbia, for which the coasts are largely MD allopatric whereas the eastern parts are sympatric, were divided in two by the 120^th^ meridian West. We extracted DNA from tissue using the Qiagen DNeasy Blood and Tissue Kit following manufacturer’s instructions and checked the samples concentration using Invitrogen Qubit assays. WGS libraries were generated at The Centre for Applied Genomics in Toronto, Canada and sequenced to an average of 4x coverage on an Illumina HiSeqX.

**Table 1:** Deer sample information

| Genome_ID | Species | Location | Sex | Group |
| --- | --- | --- | --- | --- |
| Ov_ON3 | *O_virginianus* | Ontario | F | WTD_allopatry |
| Ov_ON5 | *O_virginianus* | Ontario | M | WTD_allopatry |
| Ov_ON6 | *O_virginianus* | Ontario | M | WTD_allopatry |
| Ov_PA3 | *O_virginianus* | Pennsylvania | F | WTD_allopatry |
| Ov_VA3 | *O_virginianus* | Virginia | F | WTD_allopatry |
| Ov_AL3 | *O_virginianus* | Alabama | Unk | WTD_allopatry |
| Ov_SC1 | *O_virginianus* | South Carolina | F | WTD_allopatry |
| Ov_AB1 | *O_virginianus* | Alberta | F | WTD_sympatry |
| Ov_SK2 | *O_virginianus* | Saskatchewan | M | WTD_sympatry |
| Ov_SK3 | *O_virginianus* | Saskatchewan | M | WTD_sympatry |
| Ov_SD1 | *O_virginianus* | South Dakota | M | WTD_sympatry |
| Ov_KS1 | *O_virginianus* | Kansas | F | WTD_sympatry |
| Ov_MT1 | *O_virginianus* | Montana | M | WTD_sympatry |
| Ov_MX2 | *O_virginianus* | Mexico | M | WTD_sympatry |
| Oh_NV2 | *O_hemionus* | Nevada | M | MD_allopatry |
| Oh_CA3 | *O_hemionus* | California | Unk | MD_allopatry |
| Oh_BC6 | *O_hemionus* | British Columbia | M | MD_allopatry |
| Oh_BC8 | *O_hemionus* | British Columbia | M | MD_allopatry |
| Oh_BC1 | *O_hemionus* | British Columbia | F | MD_allopatry |
| Oh_AK1 | *O_hemionus* | Alaska | Unk | MD_allopatry |
| Oh_WA1 | *O_hemionus* | Washington | M | MD_allopatry |
| Oh_AB1 | *O_hemionus* | Alberta | F | MD_sympatry |
| Oh_UT2 | *O_hemionus* | Utah | M | MD_sympatry |
| Oh_CO1 | *O_hemionus* | Colorado | F | MD_sympatry |
| Oh_SK5 | *O_hemionus* | Saskatchewan | Unk | MD_sympatry |
| Oh_OR2 | *O_hemionus* | Oregon | Unk | MD_sympatry |
| Oh_SD2 | *O_hemionus* | South Dakota | F | MD_sympatry |
| Oh_NM2 | *O_hemionus* | New Mexico | M | MD_sympatry |

### Data processing

Raw reads quality was examined using FastQC (v0.11.9; Andrews S., 2010). We trimmed the reads using Trimmomatic (v0.36; Bolger et al., 2014), and aligned them to the WTD genome (GCA_014726795.1) with bwa-mem (v0.7.17; Li & Durbin, 2009). We sorted the reads using SAMtools sort (v1.10; Li et al., 2009), identified duplicates reads with Picard MarkDuplicates (v2.23.2; Broad Institute, GitHub Repository., 2019) and removed them using Sambamba view (v0.7.0; Tarasov et al., 2015). We performed a local re-alignment using GATK RealignerTargetCreator and IndelRealigner (v4.1.7.0; McKenna et al., 2010). For further quality checks, we used Sambamba flagstat, mosdepth (v0.3.1; Pedersen and Quinlan 2018) and MultiQC (v1.10; Ewels et al., 2016). For the MSMC analysis, some samples were sequenced to a higher depth on multiple lanes. For those, before the duplicate removal step, the read groups were specified and both sequencing files merged using Picard AddOrReplaceReadGroups and MergeSamFiles. The rest of the pipeline was performed as described above with the addition of a final step: to be able to carry out other analyses with a coverage consistent across samples, we reduced the final coverage to the average coverage of all other samples (4x) using Picard DownsampleSam.

We created three datasets for our different analyses. The first contained the variants among all deer individuals (hereafter called DeerSNP dataset). We produced it using ANGSD (v0.918; Korneliussen et al., 2014) and estimated genotype likelihoods following the GATK model (-gl 2) and called genotypes (-doGeno 4) with the following filtering: SNPs with a minimum p-value of 1e^−6^, a minimum base quality of 20 and Minimum mapQ quality of 20 (-SNP_pval 1e-6, -minMapQ 20 & -minQ 20). The second dataset additionally included invariant sites (referred to as DeerALL), it was produced following the same method as DeerSNP except for the filtering on the SNP p-value (-SNP_pval 1e-6) which was removed. Our third dataset was designed for an ABBA-BABA analysis which requires samples from an additional species as outgroup. It comprises the 28 deer individuals and one caribou sample, all mapped to the caribou genome (GCA_014898785.1; afterwards called DeerCAR dataset). For this dataset, we first sorted the deer bam files by read name and converted them to fastq using SAMtools sort and fastq respectively. We mapped these fastq to the caribou reference genome using bwa-mem. We ran ANGSD using the same procedure as for DeerSNP dataset, with the addition of a caribou genome sequence in bam format, obtained for this analysis from Dedato et al. (2022).

### Population structure analyses and cross-species coalescent rate

To infer admixture proportions and compute a PCA on allele frequencies based on genotype likelihood data we used PCAngsd (Meisner & Albrechtsen, 2018) and NGSadmix (Skotte et al., 2013) on the DeerSNP dataset. NGSadmix was run with K = 1 up to K =7 and the best K value was determined with CLUMPAK (Kopelman et al., 2015) following the Evanno method (Evanno et al., 2005) with 7 replicates for each K value. We investigated the correlation between the distribution on each PC and each individual’s range in R using the package “stats” with a linear model including the species and either latitude or longitude, and a Pearson’s correlation (e.g. lm(PC ~ Species + Latitude)).

For two samples with high coverage (Ov_ON6 at 18x - and Oh_WA1 at 22x) we implemented the multiple sequentially Markovian coalescent (MSMC) to infer the cross-coalescent rate. To do this we first generated a 35-mer mappability mask file using the SNPable pipeline as is required for MSMC (Schiffels & Wang, 2020). We restricted analyses to scaffolds >500 Kbp (Gower et al., 2018) and phased samples using whatsHap 1.3 (Martin et al., 2016) prior to generating the MSMC input files. We ran MSMC2 using the time segment pattern 1×2+25×1+1×2+1×3 and estimated the cross-coalescence rate (CCR) across populations by comparing the first haplotype from each sample in a pairwise fashion using the -P flag and skipping any sites with ambiguous phasing. To quantify the migration rate over time m(t) and the cumulative migration probability M(t) – which is related to the CCR (Schiffels & Durbin, 2014) -– we implemented the IM model to the MSMC output (Wang et al., 2020). We assumed a generation time of 2 years (Demarais et al., 2000; Deyoung et al., 2003) and mutation rate of 1.33×10-8 mutations/site/generation.

### Genome-wide ancestral introgression

To evaluate the extent of historical introgression between sympatric MD and WTD, we used the ABBA-BABA test, which allows for the detection of introgression between three populations P1, P2 & P3 and an outgroup (Durand et al., 2011; Green et al., 2010). We ran the ABBA-BABA analysis implemented in ANGSD on the DeerCAR dataset between individuals (-doAbbababa 1) and between populations (-doAbbababa2 1; Soraggi et al. 2018), using the caribou as outgroup. The D-statistic, standard error and Z-score were computed with ANGSD’s accompanying R scripts: jackKnife.R and estAvgError.R. To further our understanding of the admixture events and build a maximum likelihood tree of our system, we used treemix (v1.13; Pickrell & Pritchard, 2012) on both DeerCAR and DeerSNP datasets. We constructed the maximum likelihood trees with migration events ranging from 0 to 5, either WTD_allopatry or Caribou as root, and accounted for linkage using 1000 SNPs per block. Tree and residuals visualisations were performed with associated R script plotting_funcs.R and the variance explained by each migration event was computed with the get_f() function.

### In-windows historical introgression

The D statistic is sensitive to genomic variation (*π*), and should not be used to determine introgression on a small scale (Martin et al., 2015). The f_d_ statistic, proposed by Martin et al. (2015), is less dependent on diversity than D and therefore allows for the detection of potentially introgressed regions of the genome. We used the python script ABBABABAwindows.py (Martin, 2021) to estimate in D and f_d_ in 50Kbp windows to detect potentially introgressed loci in our DeerCAR dataset on two comparisons: 1) P1 = WTD allopatry, P2 = WTD sympatry, P3 = MD sympatry and 2) P1 = MD allopatry, P2 = MD sympatry, P3 = WTD sympatry, both with the caribou as outgroup. We then identified potentially introgressed windows as those having a f_d_ value higher than the 97.5% quantile. Their position was used in BEDTools intersect (v2.30.0; Quinlan and Hall 2010) with the caribou annotation file to identify the genes present in putatively introgressed windows.

### Genome scans

To detect islands of divergence between WTD & MD, we used the python script popgenWindows.py (Martin, 2021) to estimate individual heterozygosity, F_ST_, D_xy_ and *π* in 50Kbp windows (note we filtered out scaffolds with less than 2 windows). Since the accurate computation of D_xy_ and *π* require a dataset including the invariant sites, we performed this analysis on the DeerALL dataset. To detect the outlier loci, we based our approach on the four selection scenarios developed in (Shang et al., 2021): I) Divergence with gene flow that presents as high F_ST_ and D_xy_ but low *π*; II) allopatric selection that shows high F_ST_, low *π* and stable D_xy_; III) background selection that presents as high F_ST_ but low *π* and D_xy_; and IV) balancing selection that displays as low F_ST_ but high *π* and D_xy_. Thresholds were set as the upper or lower 5% for a high or low criteria, and between 45 and 55% for a stable condition. Outliers were flagged as belonging to one of the four scenarios when they met the specific criteria shown in Table 2.

**Table 2:** Four different evolutionary scenario and their thresholds for F_st_, *π* and D_xy_ percentiles applied in the genome scans analysis.

| Scenario | $F_{st}$ | $\pi$ | $D_{xy}$ |
| --- | --- | --- | --- |
| Divergence with gene flow (DwGF) | > 0.95 | < 0.05 | > 0.95 |
| Allopatric selection (AS) | > 0.95 | < 0.05 | Between 0.45 & 0.55 |
| Background selection (BGS) | > 0.95 | < 0.05 | < 0.05 |
| Balancing selection (BLS) | < 0.05 | > 0.95 | > 0.95 |

We identified genes present in outlier regions by comparing the coordinates of the outlier windows and the WTD annotation file in BEDTools intersect. We extracted the gene names from those regions as well as those from their associated scaffolds. The gene list from outlier windows was uploaded to ShinyGO for enrichment analysis with the scaffold’s gene list as background information and no species specification. We downloaded the enrichment analysis results for the molecular function, biological process, and cellular component GOs and analysed the results in R. We also uploaded the gene list to UniProt’s Retrieve/ID mapping tool (Pundir et al., 2016) and downloaded the information for visual inspection.

To estimate the position of WTD & MD along the speciation continuum through the correlation between recombination rate (*ρ*), F_ST_, and *π*, we computed *ρ* in windows using FastEPRR (v.2.0; Gao et al., 2016). Here, we used our four categories as a proxy for a speciation continuum with six comparisons: WTDa - WTDs and MDa - MDs representing no speciation, WTDs - MDs, WTDs - MDa and WTDa - MDs as intermediate stages and WTDa - MDa as full differentiation. We first phased the vcf using SHAPEIT (Delaneau et al., 2012) and split the resulting file into scaffolds. We then ran FastEPRR separately on our four populations to obtain the *ρ* estimation in 50kbp windows (winLength = 50000) on the 15 longest scaffolds of the WTD genome, with 100 repetitions. To be able to correlate *ρ* with F_ST_ and *π*, we re-ran the genome scan analysis on the population level. FastEPRR outputs *ρ* in 4*N*_e_*r*, to convert this value to *ρ* in cM/Mb, we computed *N*_e_ per population using the relationship *π* = 4*N*_e_μ. We used μ= 1.33×10-8 and the genome scans result to average *π* across windows in each population. After filtering to keep only windows that were present across all comparisons (12,500 windows), we generated Pearson’s correlations between I) *ρ* in both populations (*ρ*_1_ vs *ρ*_2_), II) average *ρ* between populations and F_ST_ (*ρ*_1-2_ vs F_ST_), and III) average *ρ* and average *π* between populations (*ρ*_1-2_ vs *π*_1-2_) following Shang et al. (2021). The resulting correlation coefficients were compared between all comparisons to infer trends over divergence time. All results were analysed in R version 4.1.0 “Camp Pontanezen” (R Core Team, 2021) .

## Results

We sequenced 28 individuals to an average coverage of approximately 4x. We called 103,970,889 SNPs in the DeerSNP dataset, which represented 4% of the genome. In the PCA based on allele frequencies, PC1 accounted for 59% of the variation and separated both species (Fig. 1B). PC2 explained an additional 8% of the variation and showed a spread of MD individuals that is consistent with their longitudinal distribution (Pearson correlation = 0.8; *p*-*value* = 0.00057). Latitude was not important in shaping PC1 or PC2 variation (Pearson correlation *p-value* > 0.05). Admixture analysis showed a well-defined genetic clustering between species (Evanno’s method: best K = 2, Fig. 1C, Fig. S1). The WTD individual Ov_MX2 was partially admixed in both analyses, with NGSAdmix assigning it 92.5% WTD and 7.44% MD ancestry (Fig. 1C). This individuals’ admixed pattern disappeared in the admixture analysis with higher K values (K = 3 to 7, Fig. S1), though additional within species clusters are observed. The MSMC2-IM model showed no support for migration between species once they split, with the CCR suggesting the speciation event for *Odocoileus* took place just under 1 Mya (Fig. 2).

**Figure 2.**
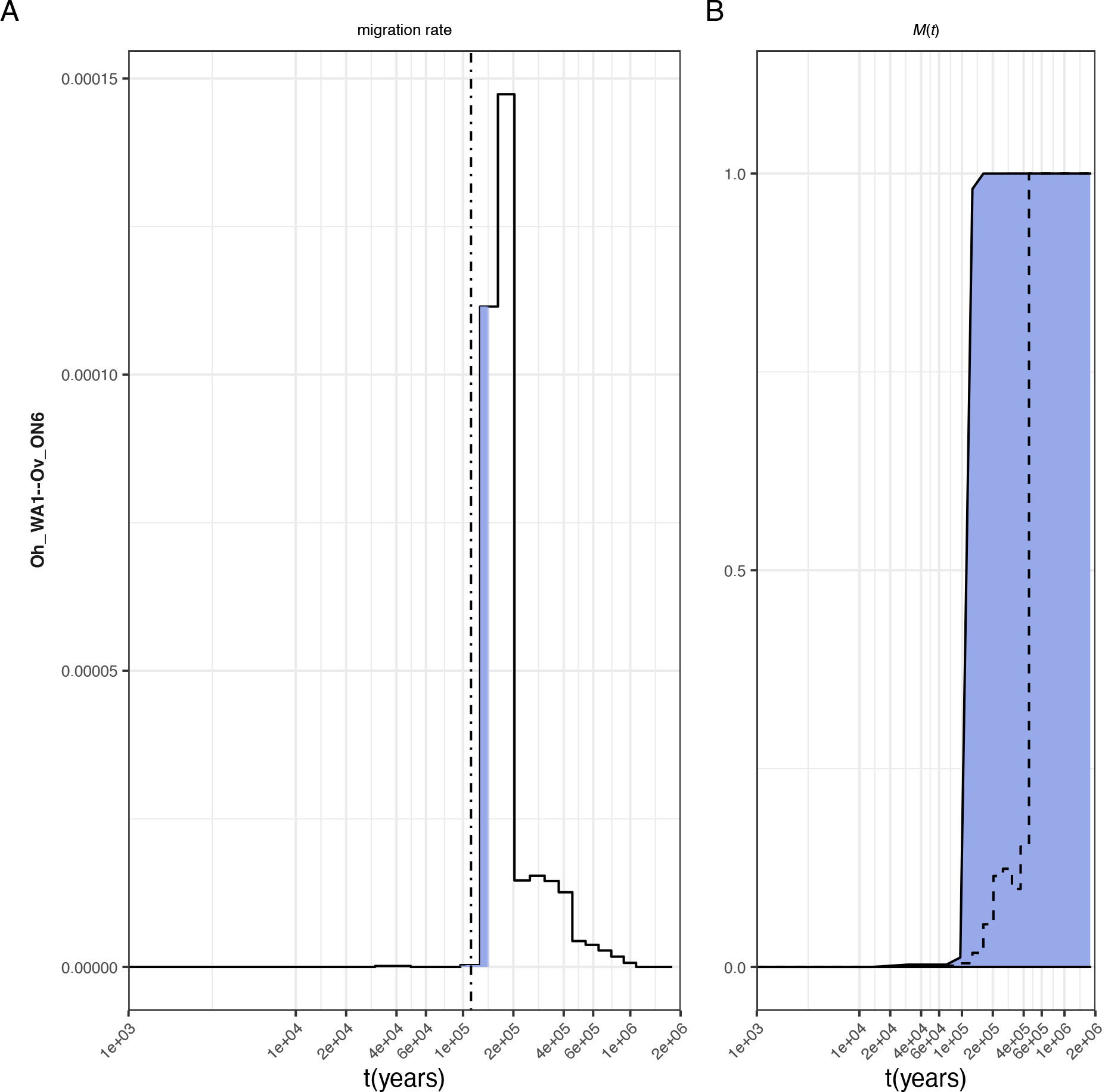
MSMC-IM analysis of white-tailed deer (Ov) and mule deer (Oh) from Ontario (ON) and Washington (WA). (A) Migration rates over time where dashed line indicates the time point where 50% of ancestry has merged. Blue shading indicates 99% percentile of the cumulative migration probability. (B) Cumulative migration probabilities *M*(*t*). Dashed lines indicate the relative cross coalescence rate obtained from MSMC2.

To detect historical gene flow, we performed an ABBA-BABA test based on genotype likelihoods between all combinations of our 28 individuals in a first analysis and between all four populations in a second, with the caribou as outgroup (DeerCAR dataset, 72,438,766 SNPs). We computed 9828 ABBA-BABA individual topologies in ANGSD, of which 30 presented signatures of introgression. We found six potential inter-species introgression events with a D-statistic ranging from 0.011 to −0.013, two of which are significant (Z-score > |3| (Barlow et al., 2021; Kirch et al., 2021), Fig. 3A). D-statistic in intra-species comparisons range from 0.25 to 0.916 in ABBA and from −0.046 to −0.914 in BABA, all intra-species comparison show a significant Z-score (Fig. 3A). In the comparisons between the four populations, three ABBA-BABA topologies presented signatures of introgression on a total of 12. The single inter-species comparison presents a D-statistic of −0.07 (Z-score = −7.66) for an introgression between sympatric MD and WTD. The other two comparisons show a reciprocal introgression between MD in allopatry and sympatry of 0.9 and 0.89 (Z-score = 1342.83 & 1376.69; Fig. S3). The treemix analyses showed a topology with no migration that explained over 99% of the variation (Fig. 3B). We further analysed historical introgression through the measure of f_d_ across 50kb non-overlapping windows in two different introgression scenarios: 1) WTDa, WTDs, MDs, Caribou and 2) MDa, MDs, WTDs, Caribou (Fig. S4). We found 1243 outlier windows with a f_d_ higher than the 97.5% percentile in the first scenario and 1248 in the second, each representing ~2.45% of the genome with 68 windows overlapping between comparisons (Fig. S4).

**Figure 3:**
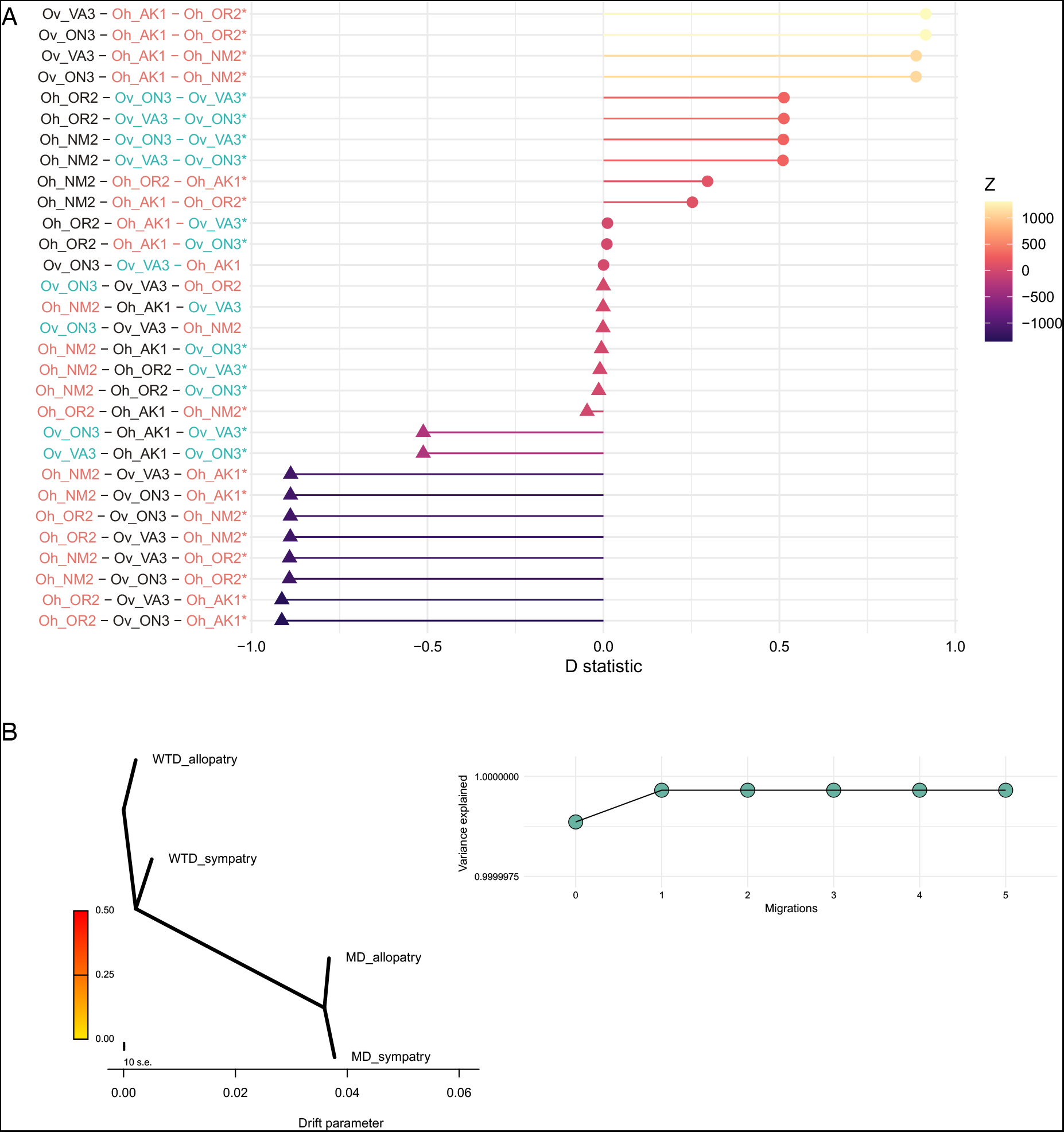
Introgression analyses. (A) ABBA-BABA analysis between individuals, individuals involved in gene flow are coloured depending on the species (WTD in blue, MD in orange), the colour gradient represents Z score, excess of ABBA depicted as points, excess of BABA shown as triangles, significant tests highlighted with an *(Z-score > |3|). (B) Maximum likelihood tree inferred by Treemix for 0 migration (right) and the variance explained by each migration event (left).

Measures of relative genomic divergence between WTD and MD were elevated (F_ST_ = 0.26) and absolute divergence d_xy_ was 0.011; genetic diversity (*π*) was higher in WTD than in MD (WTD = 0.008, MD = 0.004, Mann-Whitney test: p < 2.2e-16). We identified a total of 1183 windows presenting patterns consistent with one of the four selection scenarios (Fig. 4, Fig. S5). Of those, 1016 suggest a pattern of balancing selection, they were distributed across 236 scaffolds and represent 1.99% of the genome. Those windows contained 121 genes, some of which showed an enrichment in ontologies associated with the sense of smell and chemical stimuli, including three categories presenting an enrichment above 15-fold (Fig. S6). We also detected GOs related to the MHC and immunity, these include three categories with an enrichment above 24-fold (Fig. S6). We identified patterns of background selection in 165 windows across 58 scaffolds, representing 0.32% of the genome. These windows harboured 208 genes identified with a UniProt ID and for which we found enrichment in GOs related to epigenetic factors such as “Unmethylated CpG binding” (25-fold) or “Histone demethylation” (15-fold) (Fig. S7). These windows were either isolated or clustered together into putative islands of divergence (Fig. S5).

**Figure 4:**
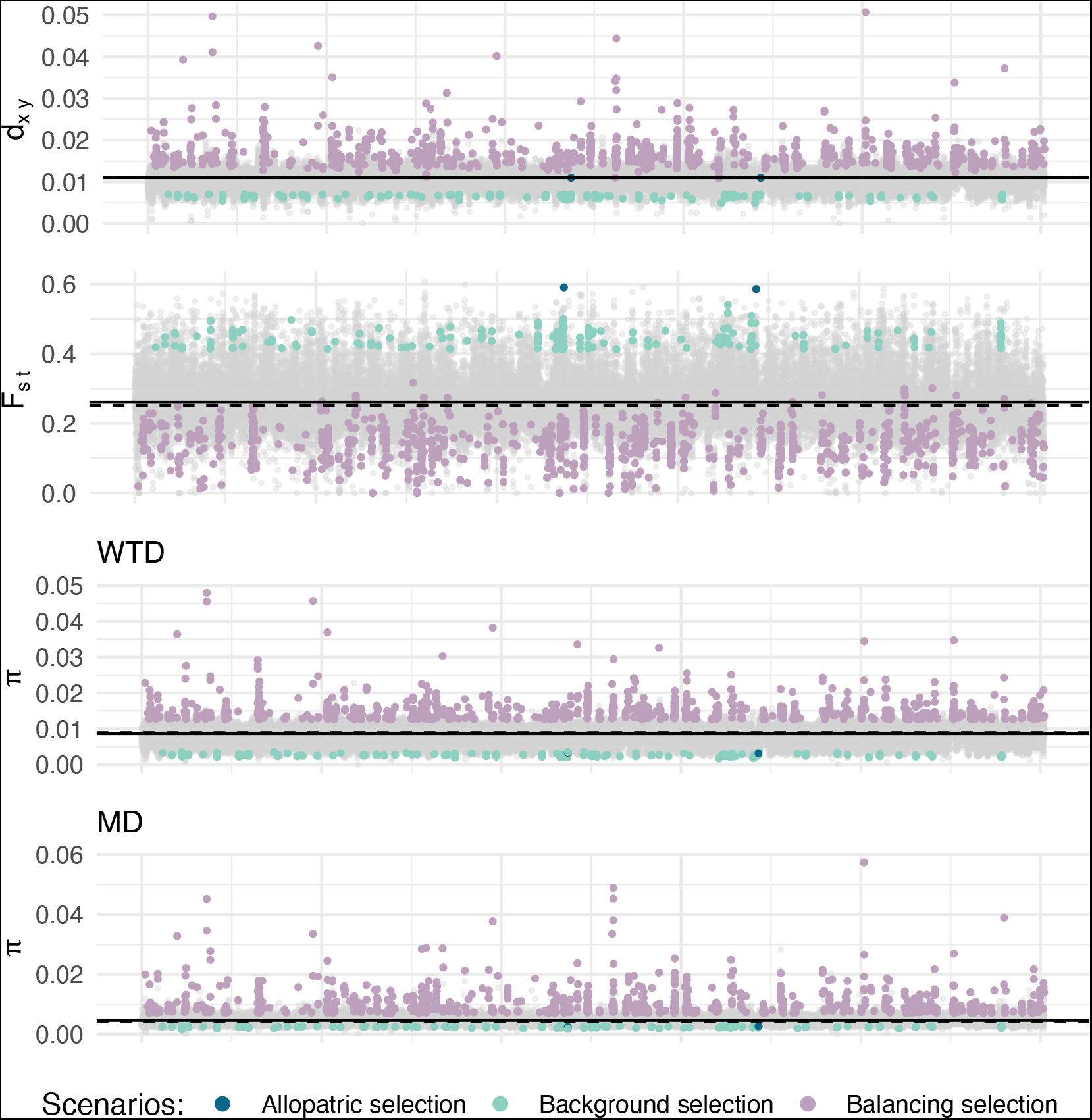
Distribution of windows presenting pattern of selection across the genome for d_xy_ (top), F_ST_ (middle) and π (bottom). Windows following a pattern of selection are coloured according to its corresponding evolutionary scenario, grey windows do not exhibit pattern of selection. The continuous and dashed line represent the mean and median respectively.

We found 2 windows under allopatric selection, each window containing one gene: ACAP2, a GTPase activating protein, and PCDHB4 potentially involved in cell-binding (UniProt, 2022a, 2022b). When we applied a more liberal cut-offs for F_ST_, d_xy_, and *π* we still observed no evidence for divergence with gene flow (Table S1). The correlations between *π*, F_ST_ and *ρ* are highly significant in all six comparisons (Fig. S8). As predicted correlations coefficients for F_ST_ - *ρ* became stronger with divergent comparisons (Fig. 5). Likewise, the *π* - *ρ* correlation coefficients were constant over divergence time, while the recombination landscape (*ρ* - *ρ*) differed between species (Fig. 5).

**Figure 5:**
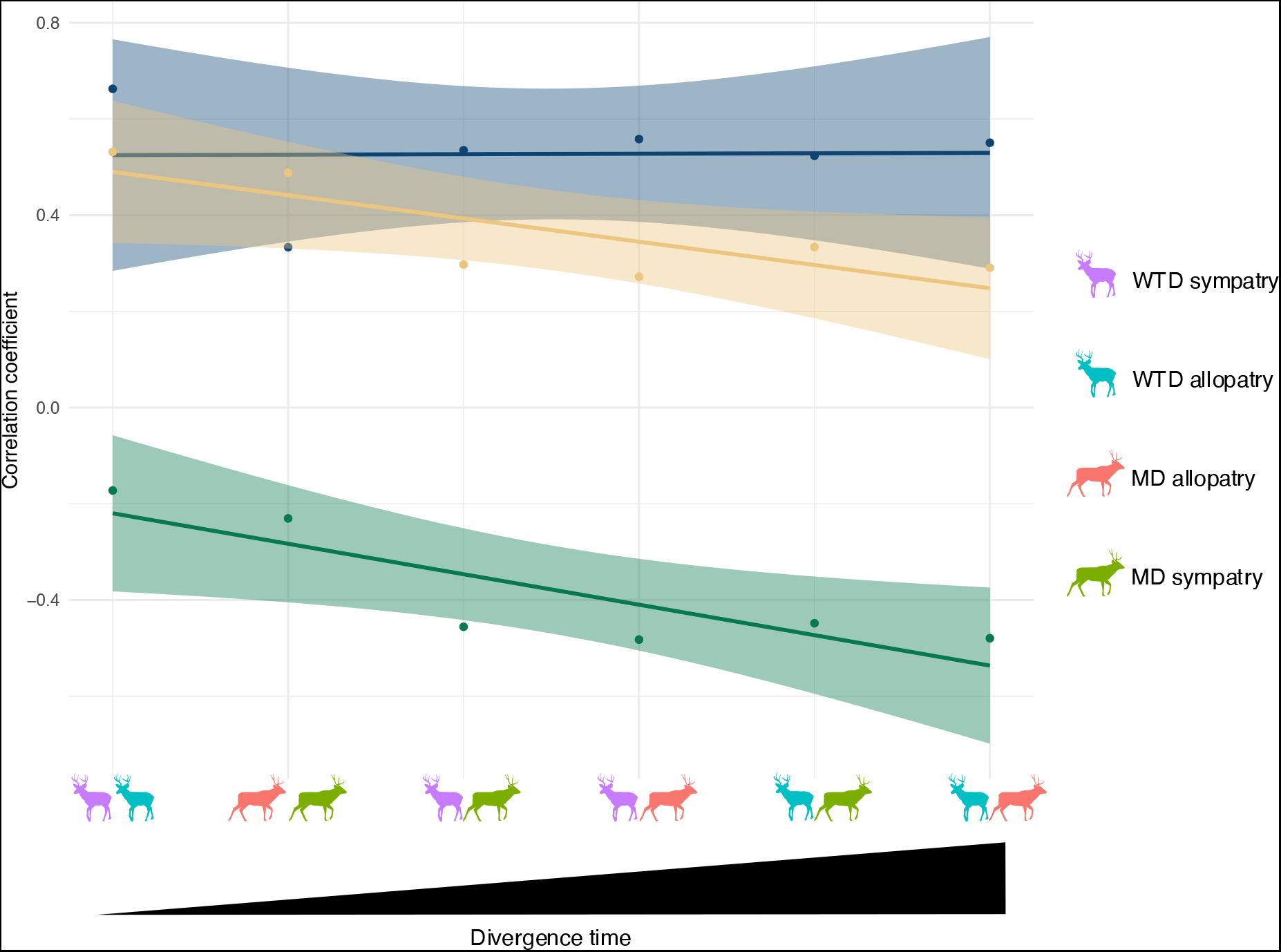
Correlation coefficients over divergence time for *π* - *ρ* (dark blue) F_ST_ - *ρ* (dark green) and *ρ* - *ρ* (yellow), trend lines represent a linear model. The speciation continuum is represented by six comparisons of our four groups).

## Discussion

We investigated the speciation history of the *Odocoileus* genera through introgression analyses and genome scans. Our results suggest speciation between WTD & MD took place with negligible historical gene flow, despite contemporary and historical evidence of interbreeding (Cathey et al., 1998; Combe et al., 2021; Cronin et al., 1988; Cronin, 1991; Derr, 1991; Hornbeck & Mahoney, 2000; Hughes & Carr, 1993; Russell et al., 2021; Stubblefield et al., 1986). Consistent with other estimates of divergence (Combe et al., 2021; Douzery & Randi, 1997), WTD and MD appear to have split ~1 Mya (Fig. 2). The main pattern of selection, however, appears to be that of balancing selection, with patterns of divergence with gene flow notably absent. Moreover, the scarcity of allopatric selection (i.e., positive selection post divergence) suggests isolation and drift primarily underlies the species differentiation, with WTD and MD likely only recently coming into secondary contact. The genome-wide scans combined with the absence of introgression suggests that WTD & MD are far along the speciation continuum (i.e. Feder 2012), despite contemporary hybridisation. Some signals of selection might have been lost by recombination, elevated *F*ST, and time, but the paucity of DwGF and AS signals even at liberal cut-offs suggest speciation was driven primarily by drift.

### Speciation with negligible historical introgression

Contemporary hybridisation in wild WTD & MD appears highly variable: rates between 1 and 19% have been observed, which can be explained by the region and degree of overlap, but also by the methodologies used that vary in resolution (Cathey et al., 1998; Combe et al., 2021; Cronin et al., 1988; Cronin, 1991; Derr, 1991; Hornbeck & Mahoney, 2000; Hughes & Carr, 1993; Russell et al., 2021; Stubblefield et al., 1986). Moreover, a proposed historical mtDNA introgression would suggest ancestral hybridisation occurred (Wright et al., 2022). We therefore expected to find signs of introgression in our samples (e.g. Combe et al. 2021) assuming that historic matings would have left some detectable signal as is seen in humans (e.g. Green et al., 2010) and other species (Liu et al., 2021; Paijmans et al., 2021; Poelstra et al., 2014). Surprisingly, we observed neglible levels of ancestral gene flow and introgression. While Combe et al. (2021) did find evidence of introgression between *Odocoileus* <u>species</u>; we note, this inference was based off a small number of topologies and included recent hybrids, both of which could skew D values. We did find signs for some contemporary admixture in one sample (Fig. 1C), but supplementary analysis with an additional WTD from Mexico showed that the admixture pattern likely reflects population structure rather than hybridisation since it is detected in both individuals (Fig. S2).

Our sampling surely is limited for the detection of recent hybridisation, as previous studies suggest gene flow is rather restricted (Cronin et al., 1988; Cronin, 1991; Derr, 1991; Hughes & Carr, 1993; Russell et al., 2021). For example, Russell et al., (2021) estimated a 1% hybridisation rate in Alberta by sampling 987 individuals in a range overlap of approximately 230,000 km^2^. While signatures of f_d_ identified potentially introgressed windows in ~1% of the genome, this metric can be a sign of incomplete lineage sorting (ILS) rather than introgression, particularly with no signs of genome-wide introgression as evidenced in the D-statistic (Durand et al., 2011). The independent treemix and MSMC analyses do not support historical gene flow, thus collectively, we consider the f_d_ signal to be reflective of ILS which would be expected given the high genome-wide diversity and a relatively recent species-split (Cutter, 2013; Combe et al. 2021).

The absence of signatures of ancestral gene flow suggest that WTD & MD evolved in allopatry and have recently come in secondary contact. The dynamic history of the North American continent, notably the glacial cycles of the Quaternary, shaped the evolution and distribution of many taxa (Avise et al., 1998; G. Hewitt, 2000; Shafer et al., 2010). The use of different refugia for prolonged periods of time during the glaciation events has increased the differentiation between populations in several species, including deer (Colella et al., 2021; Dussex et al., 2020; Ito et al., 2021; Kinoshita et al., 2020; Latch et al., 2014). Previous studies show that MD persisted in several refugia during the glacial cycles of the Pleistocene, increasing the intraspecific divergence (Latch et al., 2009, 2014; Wright et al., 2022). This divergence is highlighted here in the PCA which shows two distinct MD clusters, potentially segregating MD and its black-tailed deer subspecies (Fig. 1B). Environmental shifts following the LGM also seem to have impacted the divergence between subspecies of WTD & MD in Florida and Mexico for example (Alminas et al., 2021; Ellsworth et al., 1994). More recently, a form of allopatry was mediated through overharvest and habitat destruction of WTD where populations greatly decreased, even extirpated, in some regions (Budd et al., 2018; Chafin et al., 2021; Deyoung et al., 2003; McDonald et al., 2004). Here, the core of the current sympatric zone was greatly affected, notably Colorado, Montana, Idaho, Nebraska and Wyoming where WTD was almost extirpated before restocking efforts took place starting in the mid 40’s (McDonald et al., 2004). Historical introgression could further be hindered by selection against hybrids, either through sexual selection, intrinsic or ecological incompatibilities (Nosil, 2012; Rundle & Nosil, 2005, 2005). Given the genome-wide level of differentiation, we hypothesise that Dobzhansky-Muller incompatibles (DMIs) are in play, noting that they often occur early and rapidly (Schumer et al., 2015)

Little is known about hybrid fitness in the *Odocoileus* genus. Assessment of a single F1 male suggested it was fertile with no overt genetic defects (Derr et al., 1991). In three hybrids the spermatozoid phenotype showed a gradient of fertility with one sterile individual, one subfertile or infertile, and one potentially fertile but did not reproduce (Wishart et al., 1988); sperm phenotypic variation has been implicated in speciation (Albrechtová et al., 2012) and DMIs (Bhattacharyya et al., 2013). Sexual selection can also be a powerful driver of selection against hybrids (Servedio, 2004). In WTD, while females might favour males with larger antlers (Morina et al., 2018), sexual selection is poorly understood and most studies focus on male breeding success (DeYoung et al. 2006; DeYoung et al. 2009; Jones et al. 2011; Newbolt et al. 2017). Morphologically, however, WTD & MD are similar enough where the identification of their hybrids is often challenging with the metatarsal gland and their escape gaits being the only characters consistently distinguishing hybrids (e.g Lingle, 1992; Wishart, 1980). Thus, we propose a testable hypothesis that ecological incompatibilities combined with DMI maintain the species boundary. More research is also needed on WTD/MD hybrids to quantify their fitness and genomic regions underlying any selected hybrid phenotypes.

### Mixed signatures of selection underly speciation in North American deer

Given ongoing hybridisation between WTD & MD and a history of recurrent glacial cycles, we expected to find deer genomes showing patterns of divergence with gene flow. The absence of introgression and the nonexistence of DwGF in our genome scans, even with more liberal thresholds, is surprising (Table S1). The further absence of allopatric selection suggests a diminished role of selection in deer speciation, lending support to drift-induced model more consistent with Mayr’s (1942) view. While the number of windows in allopatric selection increased with more liberal thresholds (Table S1), they are never the dominant pattern. We suggest this absence could be a result of species being far along on the speciation continuum such that patterns of AS (specifically high F_ST_) are no longer detectable or meaningful because they were reduced with time and recombination, which is consistent with late stages of ecological speciation (Feder, 2012). Recombination rates, demography and selection can still influence the patterns investigated here. We inferred *ρ* estimates that are consistent with those in other mammals (Jensen-Seaman et al., 2004), their correlations with divergence would suggest *Odocoileus* are well defined species and reflects a shift in recombination landscape between WTD & MD (Fig. 5; (Burri, 2017; Shang et al., 2021).

Signatures of balancing selection (BLS) were widespread with windows containing genes consistent with deer biology (olfactory receptors) and previous literature (i.e., MHC genes). The major histocompatibility complex (MHC) is involved in pathogen recognition in vertebrate species (Bernatchez & Landry, 2003; Piertney & Oliver, 2006). Polymorphism in this complex is directly linked to disease susceptibility, the most diverse are the most resistant, and there is evidence that MHC diversity is maintained by BLS (Bernatchez and Landry 2003; Piertney and Oliver 2006; Aguilar et al. 2004; Pierini and Lenz 2018; Zhang et al. 2018). The same is true for the results on sensory perception and particularly the olfactory receptors genes, both of which are under BLS (Liu et al., 2021). Each allele in the olfactory receptors genes is expressed by a single sensory neuron (Degl’Innocenti & D’Errico, 2017), it is therefore plausible that diversity in these genes would be maintained by BLS. Olfactory receptors in WTD and MD, and more broadly deer, is critical to both predator detection and rutting behaviour (Ditchkoff, 2011). Anatomically, WTD has an architecture optimised for the delivery of sensory stimuli to the receptors in the nose (Ranslow et al., 2014). Chamaillé-Jammes et al. (2014) showed that MD inspect and respond differently to different predators’ olfactory cues, and both human and mammalian predators utilise wind direction and other mechanism to minimise scent detection by deer (Cherry & Barton, 2017; Zagata & Haugen, 1974). In other ungulates, olfactory reception is associated with sexual activity (Cann et al., 2019), maternal behaviour (Blank & Yang, 2017; Keller & Lévy, 2012), territory choice (Deutsch & Nefdt, 1992), predator response (Kuijper et al., 2014; Wikenros et al., 2015) and foraging (Hirata & Kusatake, 2021). Low differentiation (F_ST_) but high diversity (*π*) of olfactory receptors in deer is consistent with their biology and behaviour and would be expected to be under strong selection pressures.

## Conclusion

Our results suggests that WTD & MD do not conform to a speciation with gene flow scenario despite evidence of contemporary hybridisation. We propose they were spatially separated during the Quaternary glaciation cycles where genome-wide differentiation accrued via drift. This is evidenced by the majority of the genome (> 97%) not matching a selection scenario. Increased sampling and model-based demographic assessment should help clarify the role of glaciers and secondary contact in North American deer. The near absence of patterns of allopatry and ancestral gene flow suggest that white-tailed and mule deer are far along the speciation continuum, the absence of introgression signs could suggest DMIs and selection against hybrids which would contribute to the reinforcement of reproductive isolation. Future studies should focus on assaying hybrid phenotypes and vigour.

## Acknowledgements

Camille Kessler was supported by an International Graduate Scholarship. Eric Wootton was supported by three Natural Sciences and Engineering Research Council (NSERC) Undergraduate Student Research Awards. This work was supported by NSERC Discovery Grant (Grant Number: RGPIN-2017-03934); ComputeCanada Resources for Research Groups (Grant Number: RRG gme-665-ab); Canadian Foundation for Innovation: John R. Evans Leaders Fund and the Ontario Early Researcher Award (Grant Number: #36905). We thank Catherine Cullingham, Anh Dao, Orrin Duvuvuei, Brad Fulk, Steve Griffin, Levi Jaster, Lee Jeffers, Emily Latch, the NRDPFC, Charles Ruth, David Walter, Geoff Williams, Kevin White, Mark Wong, Kiana Young and Liana Zanette, for providing samples. We are also grateful to Andrew Foote, Jose Alberto Lòpez Alemàn and Emily Latch for their comments on the manuscript. Trent University is located on the traditional territory of the Michi Saagiig Anishnaabeg. Our sampling also covers most of North America, a stolen land which remains a home to many First Peoples. We are grateful to have had the opportunity to work on this land, would like to show our respect to the First Peoples, and thank them for their care, stewardship, and teachings. Miigwetch.

## Data Accessibility and Benefit-Sharing

### Data Accessibility

Raw reads for the 28 deer individuals were deposited on the NCBI Sequence Read Archive (Accession number PRJNA830519). All scripts are available on GitLab (https://gitlab.com/WiDGeT_TrentU/graduate_theses/-/tree/master/Kessler/CH_01).

### Benefit-Sharing

The contributions of tissue samples to the research are described in the ACKNOWLEDGEMENTS, benefits from this research results in data accessible on public databases as described above.

### Author contributions

CK and ABAS conceived the study, CK performed bioinformatic analyses with contribution from ABAS, CK and EW performed the molecular laboratory work, CK and ABAS wrote the manuscript. All authors commented on the text.

## Notes

https://gitlab.com/WiDGeT_TrentU/graduate_theses/-/tree/master/Kessler/CH_01

### Competing Interest Statement

The authors have declared no competing interest.

### Summary of Updates

Manuscript accepted for publication, only minor changes from the previous version

