## Supplementary Figures and tables for "Speciation without gene-flow in hybridising deer"

### Supplemental information

**Table S1:** Number of windows in the evolutionary scenarios using different thresholds, (†) indicates which thresholds we used for the analyses.

| Thresholds | DwGF | AS | BGS | BLS | Total |
|---|---|---|---|---|---|
| † Lower = 5%<br>Upper = 95%<br>Stable = Between 45 & 55% | 0 | 2 | 165 | 1016 | 1183 |
| Lower = 10%<br>Upper = 90%<br>Stable = Between 35 & 65% | 0 | 35 | 961 | 2314 | 3310 |
| Lower = 15%<br>Upper = 85%<br>Stable = Between 30 & 70% | 0 | 153 | 2007 | 3658 | 5818 |
| Lower = 20%<br>Upper = 80%<br>Stable = Between 20 & 80% | 2 | 739 | 2977 | 5207 | 8925 |

Table S2: Additional sample information, approximate sampling coordinates.

| Genome_ID | Location | Latitude | Longitude |
|---|---|---|---|
| Ov_ON3 | Ontario | 42.4800 | -82.0000 |
| Ov_ON5 | Ontario | 46.3600 | -84.0000 |
| Ov_ON6 | Ontario | 44.5300 | -78.0500 |
| Ov_PA3 | Pennsylvania | 39.8944 | -80.0870 |
| Ov_VA3 | Virginia | 38.1300 | -78.1600 |
| Ov_AL3 | Alabama | 32.3823 | -85.6858 |
| Ov_SC1 | South Carolina | 33.4148 | -80.4105 |
| Ov_AB1 | Alberta | 52.7800 | -111.0148 |
| Ov_SK2 | Saskatchewan | 52.8600 | -102.3600 |
| Ov_SK3 | Saskatchewan | 53.6330 | -109.2000 |
| Ov_SD1 | South Dakota | 44.4415 | -102.6855 |
| Ov_KS1 | Kansas | 38.6900 | -100.8158 |
| Ov_MT1 | Montana | 47.5275 | -113.7110 |
| Ov_MX2 | Mexico | 30.7295 | -108.7348 |
| Oh_NV2 | Nevada | 41.3036 | -115.1377 |
| Oh_CA3 | California | 35.3733 | -119.0187 |
| Oh_BC6 | British Columbia | 54.7824 | -127.1686 |
| Oh_BC8 | British Columbia | 49.1579 | -121.9515 |
| Oh_BC1 | British Columbia | 48.6955 | -123.3223 |
| Oh_AK1 | Alaska | 57.6853 | -134.4895 |
| Oh_WA1 | Washington | 46.8806 | -122.4092 |
| Oh_AB1 | Alberta | 52.6251 | -110.7518 |
| Oh_UT2 | Utah | 40.9733 | -111.6903 |
| Oh_CO1 | Colorado | 39.4853 | -108.0932 |
| Oh_SK5 | Saskatchewan | 50.7442 | -107.8104 |
| Oh_OR2 | Oregon | 44.4100 | -122.5340 |
| Oh_SD2 | South Dakota | 43.2240 | -103.4512 |
| Oh_NM2 | New Mexico | 36.1846 | -103.5916 |

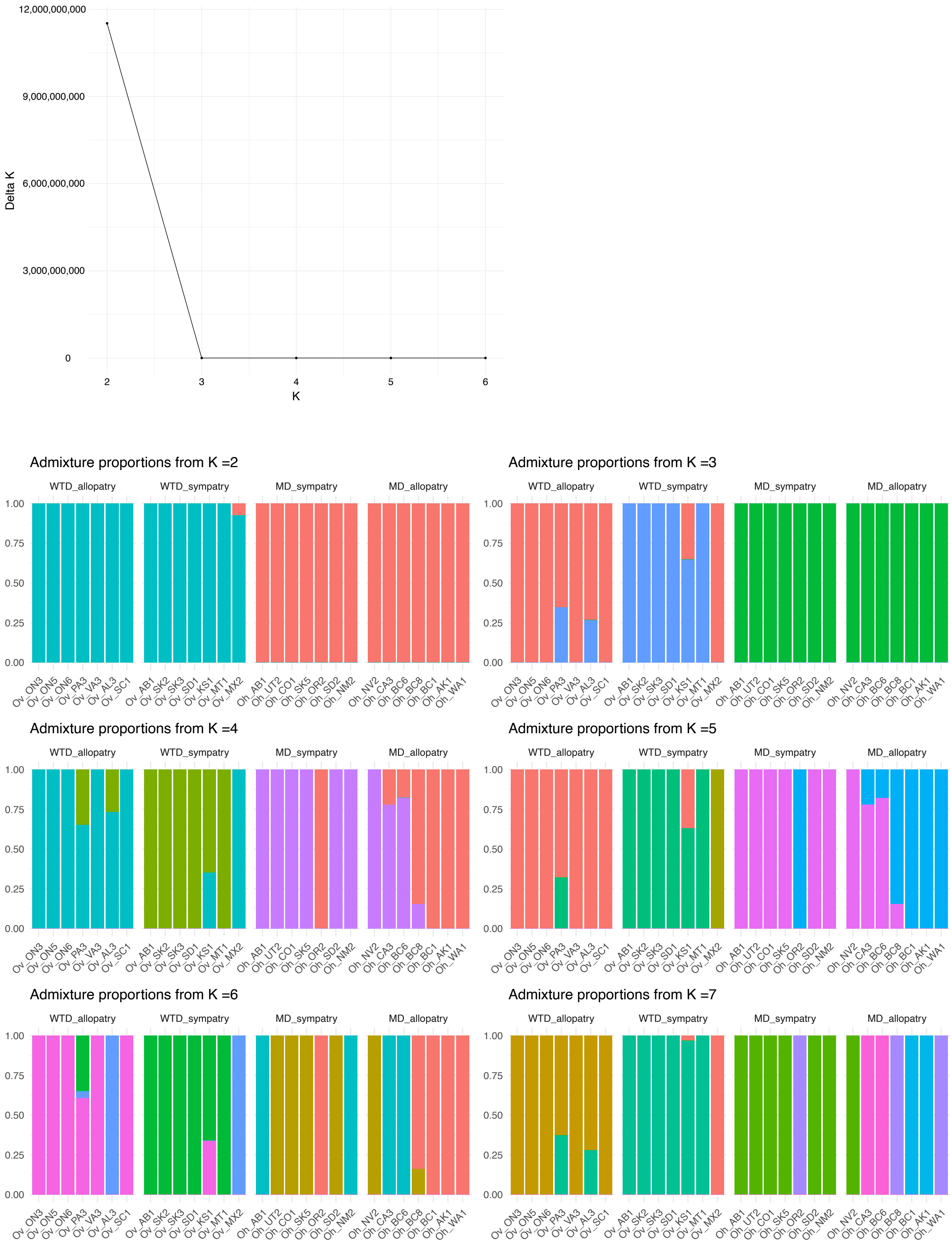


Figure S1: Evanno plot for K = 2 through to  K = 6 (top). Estimates of admixture proportions from NGSAdmix for different values of K, from K = 2 through K = 7 (bottom).

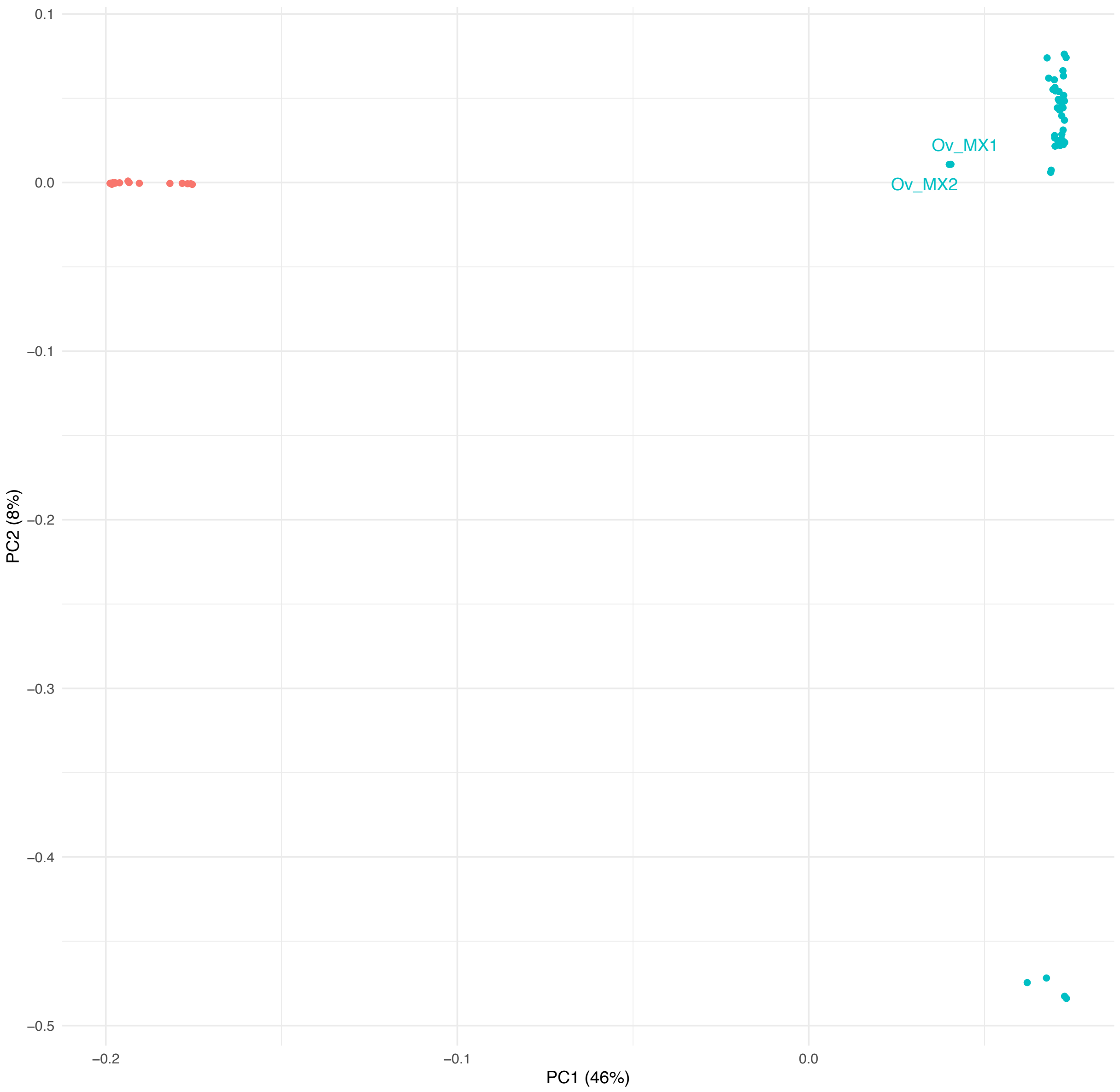


Figure S2: PCA of 73 *Odocoileus* individuals from unpublished data. WTD coloured un blue, MD in orange.

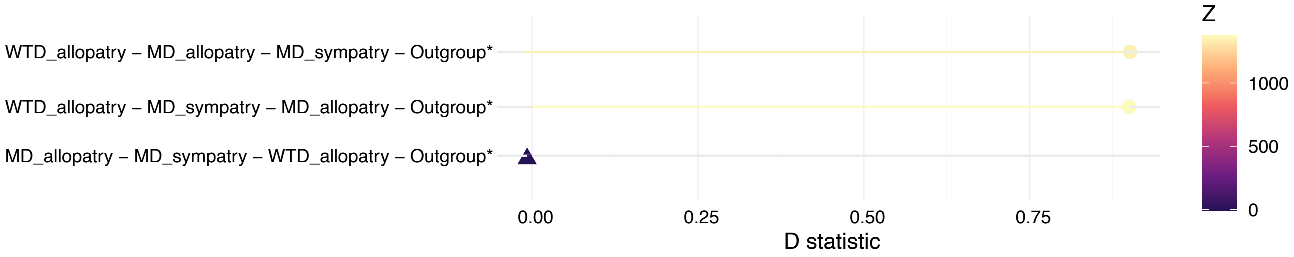


Figure S3: ABBA-BABA analysis between populations, colour gradient represents Z-score, excess of ABBA depicted as points, excess of BABA shown as triangles.

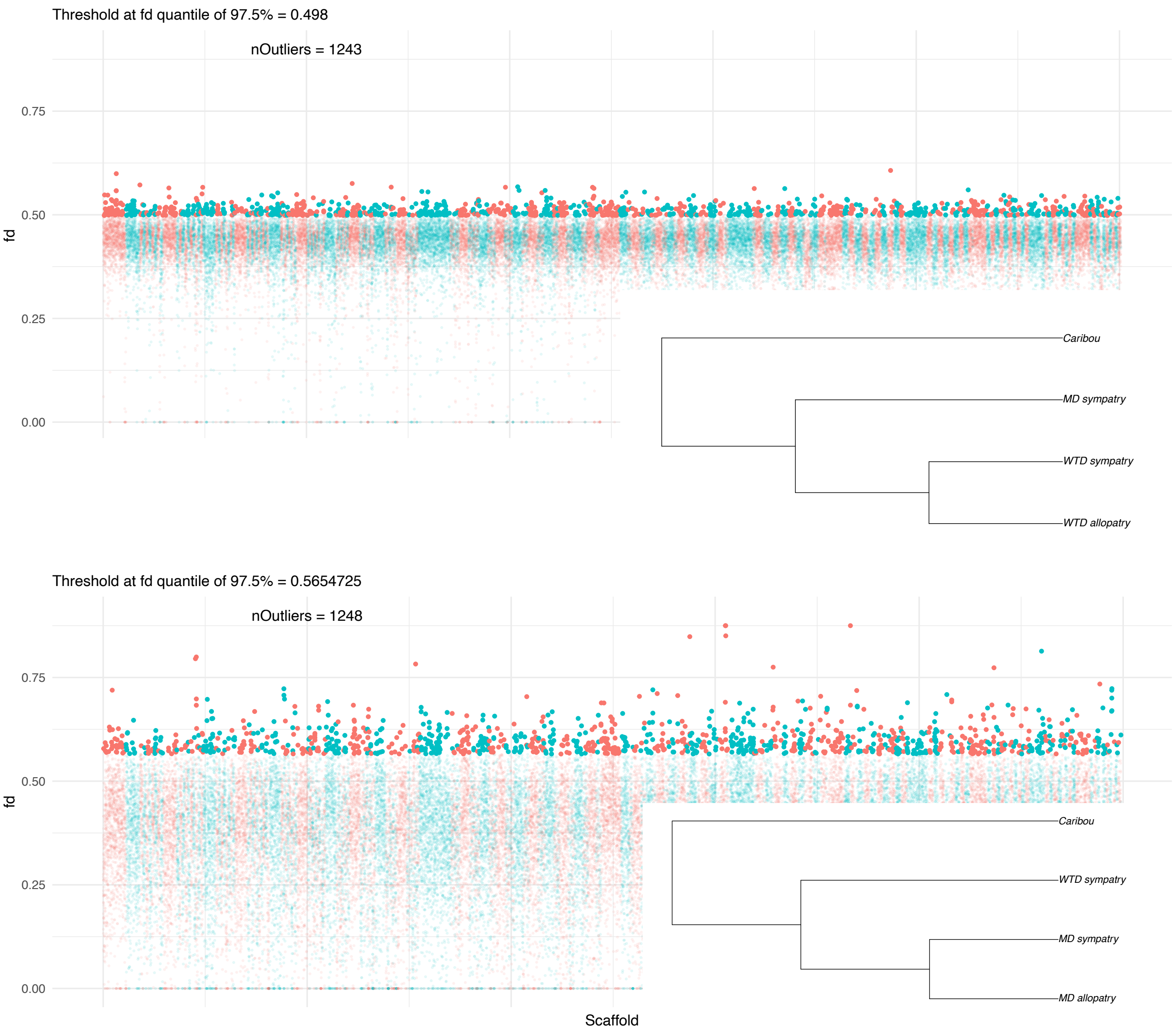


Figure S4: In windows ABBA-BABA outliers of fd for two comparisons

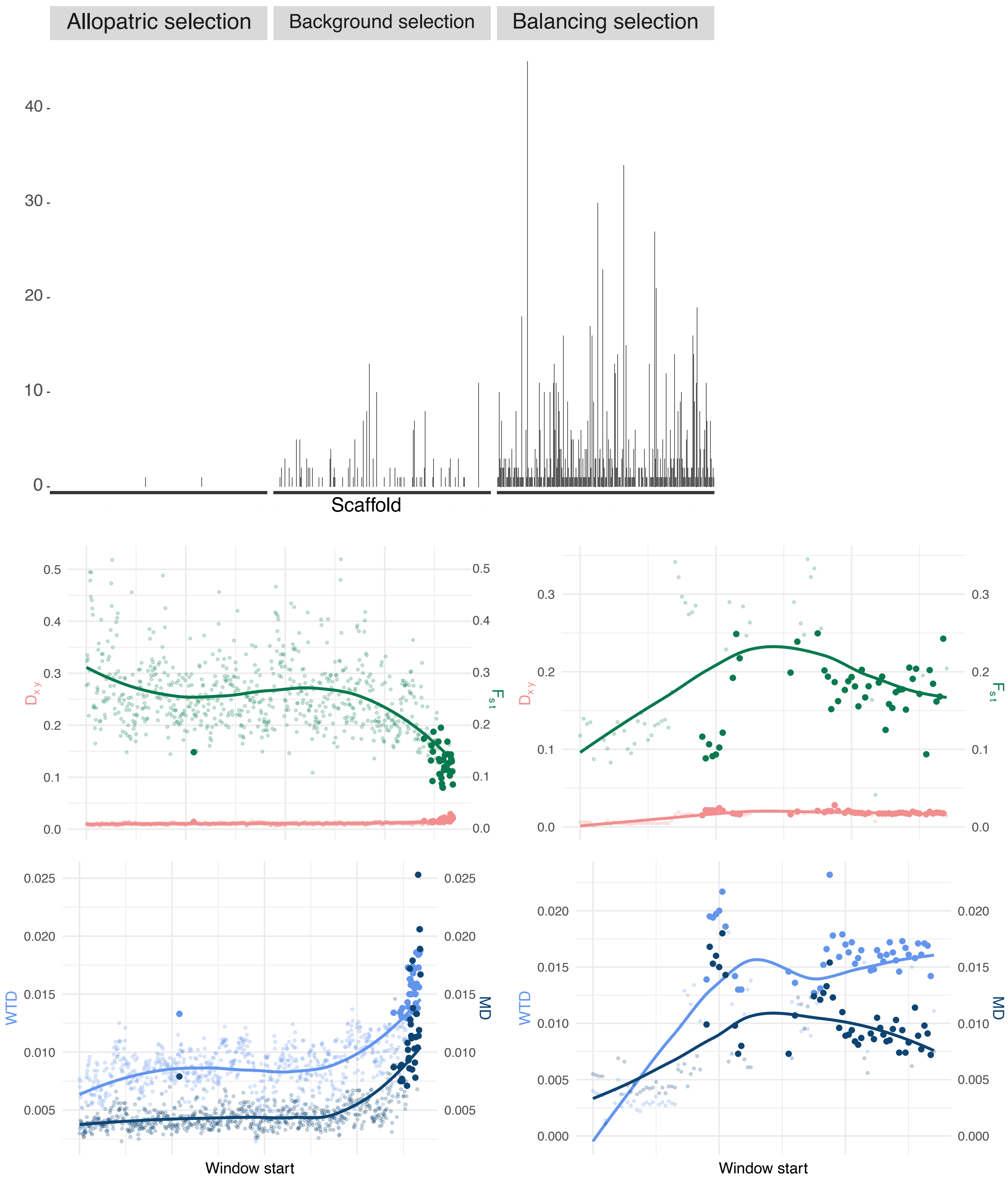


Figure S5: Count of outlier windows per scaffolds (top) and example of outliers in balancing selection across scaffold 1816 (right) and scaffold 393 (left; Outlier in darker shades)

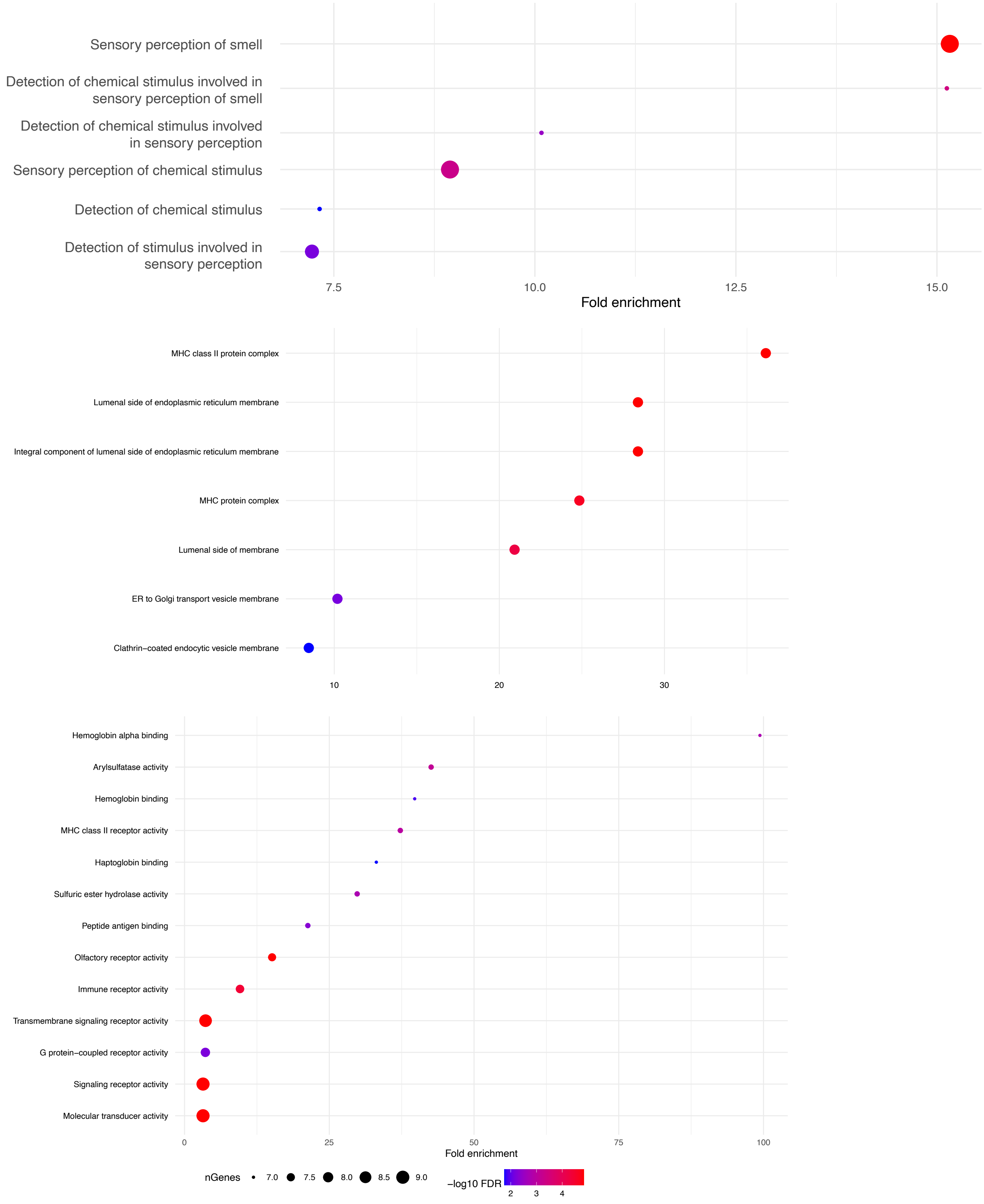


Figure S6: Gene enrichment analysis for windows in balancing selection for three GO categories: Biological Process (top), Cellular Component (middle) and Molecular Function (bottom)

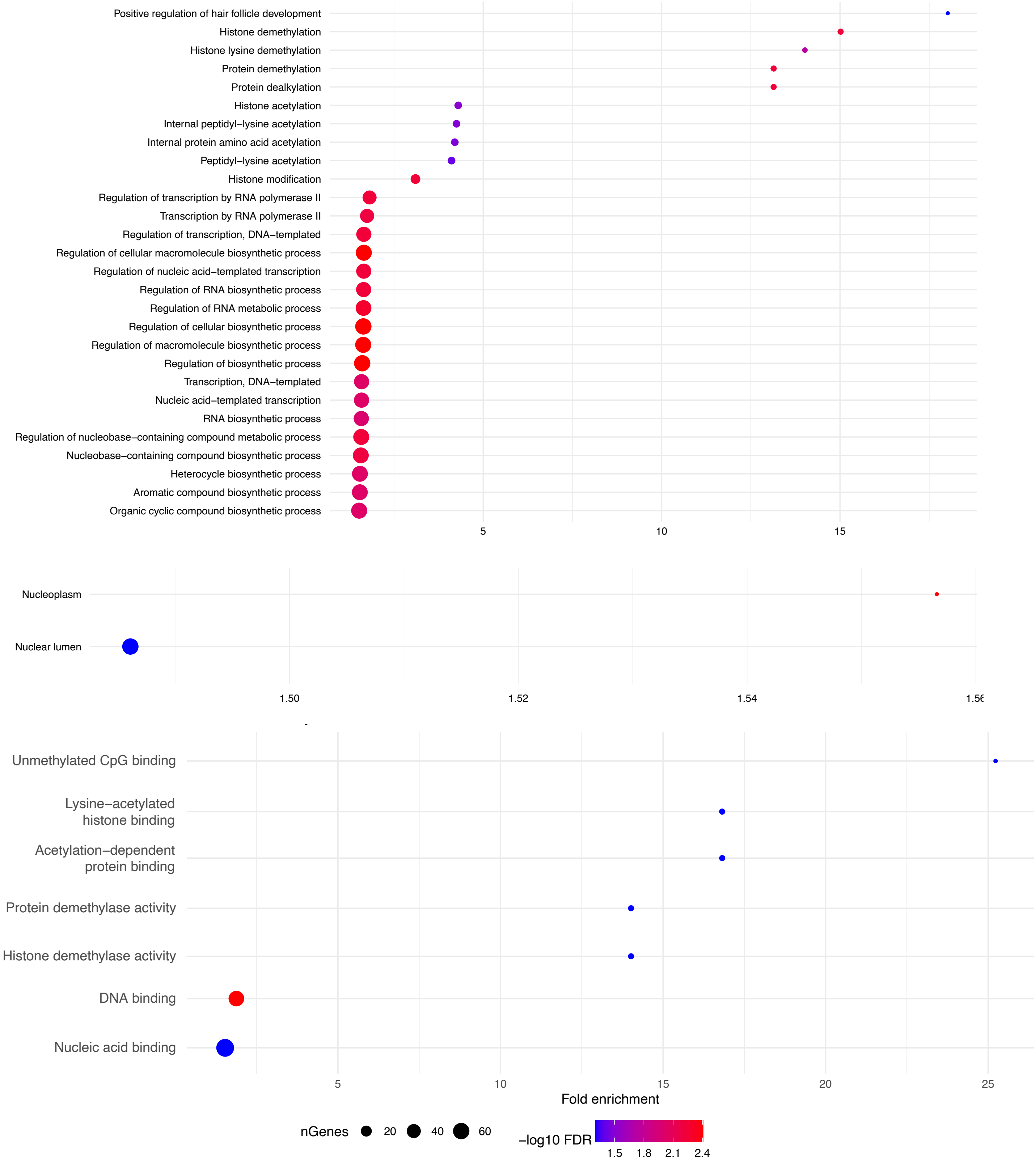


Figure S7: Gene enrichment analysis for windows in background selection for three GO categories: Biological Process (top), Cellular Component (middle) and Molecular Function (bottom)

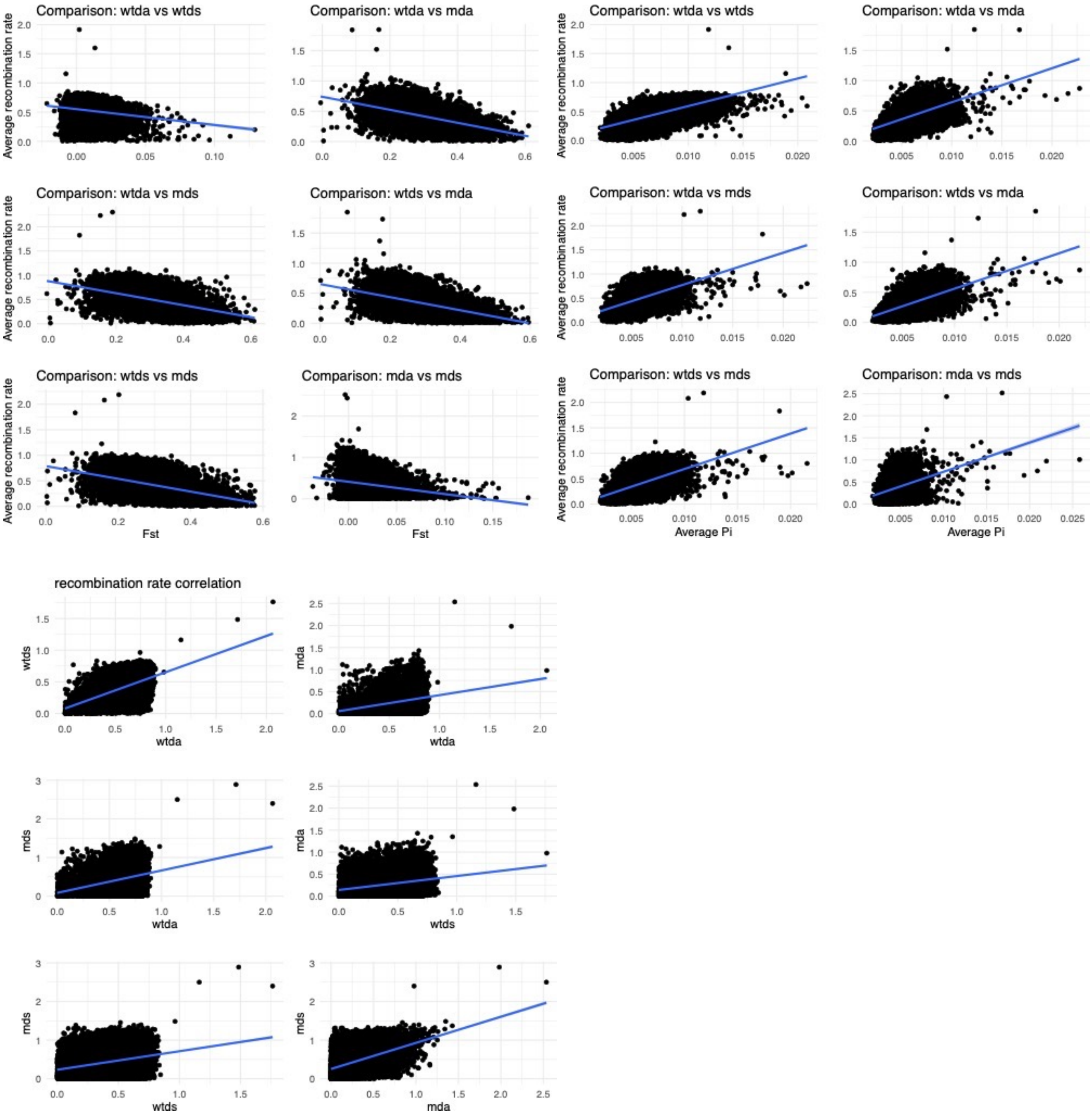


Figure S8: Correlations between $\rho_{1-2}$ vs $F_{ST}$ (top-left), $\rho_{1-2}$ vs $\pi_{1-2}$ (top-right) and $\rho_1$ vs $\rho_2$ (bottom-left) for all six comparisons
